# Probe set selection for targeted spatial transcriptomics

**DOI:** 10.1101/2022.08.16.504115

**Authors:** Louis B. Kuemmerle, Malte D. Luecken, Alexandra B. Firsova, Lisa Barros de Andrade e Sousa, Lena Straßer, Lukas Heumos, Ilhem Isra Mekki, Krishnaa T. Mahbubani, Alexandros Sountoulidis, Tamás Balassa, Ferenc Kovacs, Peter Horvath, Marie Piraud, Ali Ertürk, Christos Samakovlis, Fabian J. Theis

## Abstract

Targeted spatial transcriptomics methods capture the topology of cell types and states in tissues at single cell- and subcellular resolution by measuring the expression of a predefined set of genes. The selection of an optimal set of probed genes is crucial for capturing and interpreting the spatial signals present in a tissue. However, current selections often rely on marker genes, precluding them from detecting continuous spatial signals or novel states. We present Spapros, an end-to-end probe set selection pipeline that optimizes both probe set specificity for cell type identification and within-cell-type expression variation to resolve spatially distinct populations while taking into account prior knowledge, as well as probe design and expression constraints. To facilitate data analysis and interpretation, Spapros also provides rules for cell type identification. We evaluated Spapros by selecting probes on 6 different data sets and built an evaluation pipeline with 12 quality metrics to find that Spapros outperforms other selection approaches in both cell type recovery and recovering expression variation beyond cell types. Furthermore, we used Spapros to design a SCRINSHOT experiment of adult lung tissue to demonstrate how probes selected with Spapros identify cell types of interest and detect spatial variation even within cell types. Spapros enables optimal probe set selection, probe set evaluation, and probe design, as a freely available Python package.

## Introduction

Single cell transcriptomics has enabled the study of tissue heterogeneity at an unprecedented scale and resolution^1,2^. Recently, spatial transcriptomics (ST) technologies have added spatial context to these measurements to describe both tissue composition and organization^3,4,5,6^. Yet, this additional information requires a compromise either in spatial resolution or in the number of measured features. While untargeted ST methods aggregate measurements over multiple cells and thus lack single-cell resolution^7–9^, targeted ST methods measure the expression of a limited number of genes^10–15^. Selecting which genes to target is crucial for successful targeted ST experiments.

Probe set selection must be guided by analysis goals and the limitations of the experimental technique. Typical analysis goals include the identification of cell types, the description of cell states and transitions, and the spatial characterization of cell communication patterns and active multicellular programs. Thus, any selected gene set needs to include cell type marker genes while also capturing general transcriptional heterogeneity beyond cell type annotations. Simultaneously, one must account for technical limitations on the expression levels (e.g., due to optical crowding^16,17^), and constraints on probe design, and allow users to include prior knowledge such as pre-selected genes that may be relevant to disease studies.

Selecting probe sets for targeted ST experiments is a feature selection problem. Typically, expression profiles from dissociated cells are used as reference^4,18^ (Fig 1a). A central assumption is that genes that show interesting transcriptional variation in dissociated data will show the same in targeted ST experiments: cell type markers and highly variable genes from single cell RNA sequencing (scRNA-seq) data should highlight cell types and genes with interesting spatial patterns. Feature selection applications frequently used in scRNA-seq data analysis include highly variable gene selection, marker gene detection, and module detection^19^. Several of these approaches have also been applied to probe set selection^3,4,6^. Dedicated probe set selection approaches that use a reference scRNA-seq dataset to optimize for cell type classification^20–30^, or the capture of transcriptional variation^31–34^ have also been proposed. However, few approaches account for both cell-type and gene variation, and none of the above methods include technical constraints in their selection procedure. Since the number of genes that can be selected is limited, probe set selection is a combinatorial problem: an optimal probe set consists of those genes that together optimize multiple objectives simultaneously. Separating probe selection from probe design (Fig 1b) neglects the combinatorial nature of the problem. Additionally most available methods are non-combinatorial, score based methods and therefore rather helper-tools for laborious manual selections. To tackle the probe set selection problem as a whole, combinatorial selection is required, ideally providing interpretable combinatorial rules for practical downstream analysis.

**Fig. 1:**
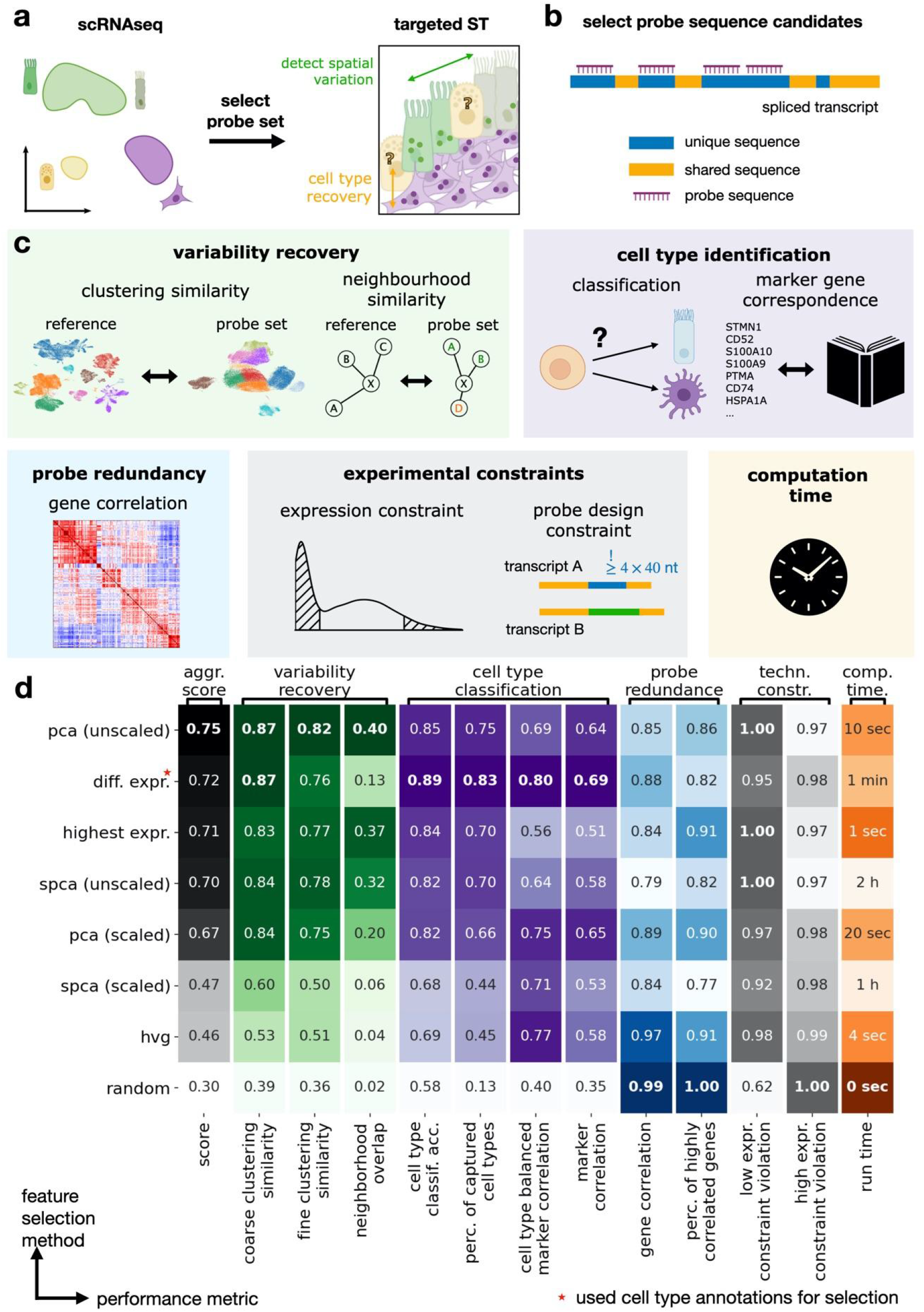
Probe set selection problem and evaluation of selected probe sets. **a**, Schematic of the probe set selection problem. A gene set is selected on scRNA-seq data and used for targeted spatial transcriptomics. The gene set is optimized to identify cell types of interest and to capture cellular variation beyond cell types. **b**, Schematic of the probe design constraint. To measure a specific gene’s expression there must be enough unique probes that can be designed. The unique sequences only occur in at least the expressed isoforms of the targeted gene and not in any RNA of other genes. Sequences that do not have that property are labeled as shared. **c**, Schematic diagram of our test suite to evaluate suitability of selected gene sets for targeted spatial transcriptomics experiments. The test suite includes multiple metrics that are categorized in variation recovery, cell type classification, gene redundancy, computation time, fulfillment of experimental constraints. **d**, Performance comparison for gene sets selected with basic feature selection methods. The aggregated score is the average between variation recovery metrics and the first two cell type classification metrics.

Here we present Spapros, a combinatorial probe set selection pipeline that takes into account prior knowledge, technical constraints, and probe design, while optimizing simultaneously for cell type identification and transcriptional variation. To evaluate Spapros, we developed a suite of evaluation metrics that measure transcriptional variation recovery, cell type identification, redundancy of genes, and fulfillment of technical constraints. In our benchmark Spapros outperforms other methods in both cell type identification and variation recovery. Using Spapros to design a probe set for a SCRINSHOT^13^ experiment of human adult lung tissue, we show that Spapros probes identify cell types of interest and detect spatially relevant variation also between cells of the same type. Spapros enables optimal experimental design for targeted spatial transcriptomics and rapid comparison of proposed probe sets through our user-friendly evaluation suite.

## Results

### Quantifying optimal probe set selection

Optimal experimental design can be guided and evaluated by quantifying how suitable candidate probe sets are for downstream analysis of the spatial data. As it is infeasible to perform a new spatial experiment for each proposed probe set, we must find proxies for exploratory analysis success. These proxies should include both the ability to identify known biology (cell type identification) and represent cellular variation that may be found (variation recovery). Following these typical analysis goals, we developed 12 metrics to measure performance in these orthogonal categories, as well as probe redundancy, violation of technical expression constraints, and selection run time (see **Methods**; Fig 1c).

Cell type identification metrics measure cell type *classification accuracy* and the *percentage of captured cell types* (Fig. S1a), and additionally how well the marker expressions of a literature derived list are captured via *marker correlation* and *cell type balanced marker correlation* (Fig. S1b). Variation recovery metrics measure how well cellular variation of the full transcriptome is recovered with only a subset of features. These comprise c*oarse* and *fine clustering similarity*, which quantify how well the gene set recovers cluster structure at different levels of granularity, and n*eighborhood similarity, which* measures how well the local cell neighborhoods are preserved (Fig. S2). The amount of redundant genes in the gene set is assessed via *gene correlation* and the *percentage of highly correlated genes* (Fig. S3). The *Low* and *high expression constraint violation* metrics measure how strongly the gene set violates technical expression thresholds. Finally, we measure the *computation time* of the feature selection methods. The overall performance of a probe set is then computed as the average of the variation recovery metrics and the cell type identification metrics *classification accuracy* and *percentage of captured cell types* as these are the main objectives we want to optimize for. We integrated these metrics into a modular, reproducible Nextflow^35^ pipeline for probe set evaluation using our Spapros Python package. The Spapros evaluation pipeline is freely available at https://github.com/theislab/spapros-pipeline. It enables large scale evaluations by automated parallelised HPC usage, and can be used to compare probe set selection methods as well as manually selected probe sets.

### Classical feature selection methods optimize different probe set selection objectives

Feature selection approaches are widely used in typical scRNA-seq data analysis pipelines^19^. As such, we investigated whether these approaches are also suitable for probe set selection. We applied our evaluation suite to investigate the performance of several general feature selection methods (based on PCA, sparse PCA (SPCA), differential expression (DE) and highly variable genes (HVG); see **Methods**). Additionally, we added random selections and a set of highest expressed genes as baseline comparisons (Fig 1d and S1-S3).

Overall, feature selection methods perform well at particular probe set selection objectives, but no method outperforms others across all metrics. For example, PCA-based feature selection (on unscaled data) clearly outperforms other methods in variation recovery aspects. This difference is most evident when considering finer cellular substructure (*fine clustering similarity* and *neighborhood similarity*, which measure recovery of cell state variation), and to be expected given PCA’s aim to reconstruct maximal variation. For recovery of coarse effects (coarse clustering similarity) we find similar performance for PCA and DE feature selections. Interestingly, the set of highest expressed genes ranks second in variation recovery and PCA-based selection on unscaled data, the top performer in this category, also introduces a natural bias towards highly expressed genes. Considering the performance of highest expressed genes and PCA-based selection on scaled vs unscaled data, this bias appears to be beneficial for variation recovery in scRNA-seq data (Fig. S2). For cell type identification, differentially expressed genes score highest among basic feature selection methods. Since PCA-based selection also ranks highly on cell type identification we observe that optimisation for variation recovery and cell type identification can go hand in hand.

As expected, random gene selections exhibit the lowest probe redundancy. Yet, optimizing this metric may not be desirable as few correlated genes can increase robustness to noise, and medium levels of correlation (~0.3 - 0.6) between two genes do not preclude that important information is gained by selecting both. In contrast, high levels of correlation (as exhibited by the SPCA-based selection, Fig. S3) leads to lower information content in the limited probe set. A balance appears to be struck by the best performing methods (PCA, DE).

Finally, basic feature selection methods do not take into account technical constraints such as gene expression limits due to image saturation or optical crowding and probe design limitations. The expression constraint metrics show that technical constraints are violated by simple feature selection approaches.

Overall, feature selection methods using DE genes or PCA perform well at individual aspects of probe set selection, but no method addresses all objectives at once. Thus, these methods are well-suited as components of a larger probe set selection pipeline, which must additionally account for technical constraints.

### End-to-end probe set selection with Spapros

Based on the results of our feature selection benchmark, we built the Spapros pipeline: an end-to-end probe set selection pipeline using PCA-based and DE gene selection as a basis (Fig 2a). The Spapros pipeline performs optimized gene selection and designs the probe sequence while accounting for technology-specific technical constraints. These aspects are considered jointly to deliver an optimal combinatorial probe set that can directly be ordered without the need for further probe filtering.

**Fig. 2:**
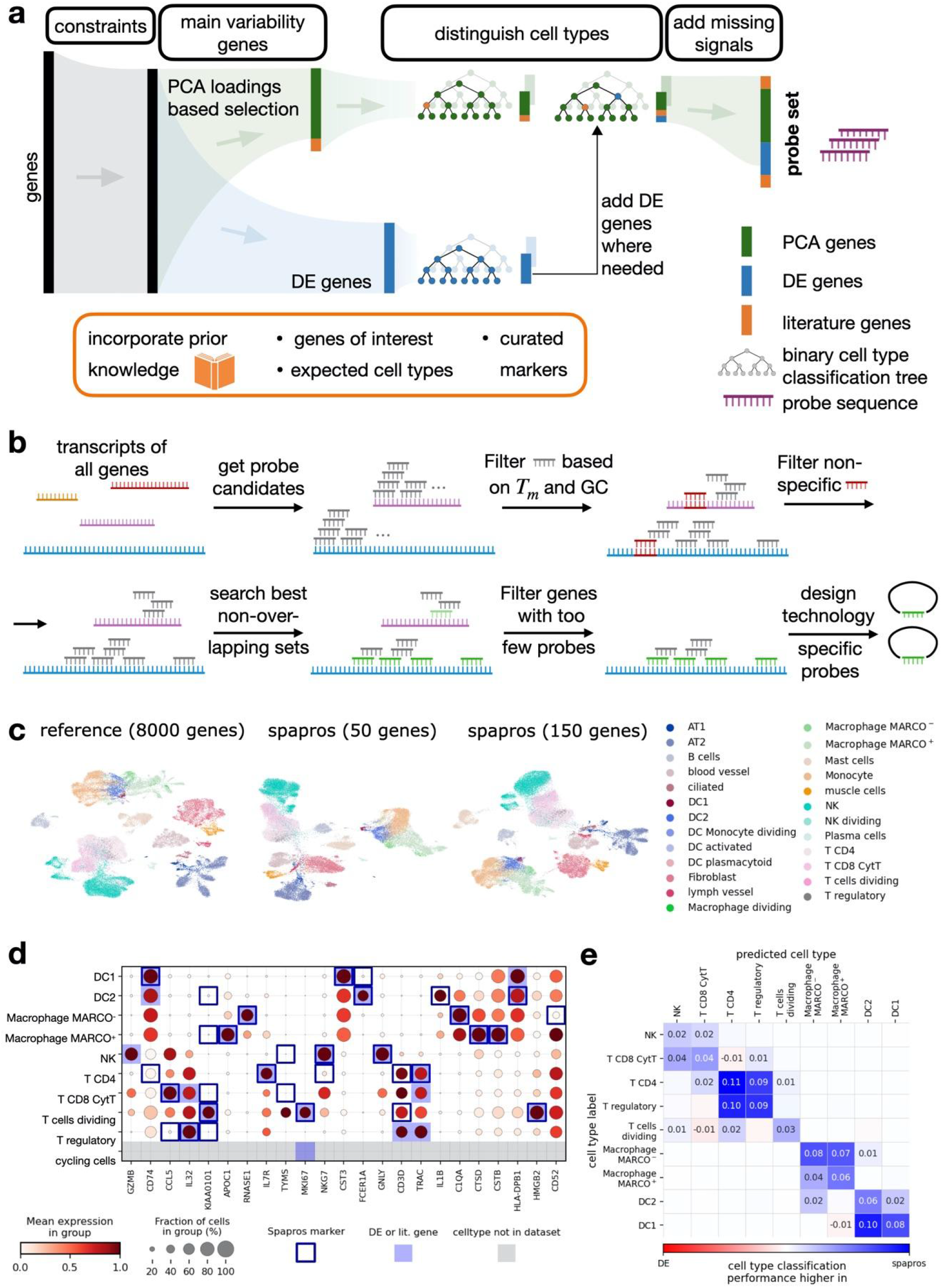
The Spapros probe set selection pipeline. **a**, Schematic diagram of the probe set selection pipeline. After pre-filtering genes on probe design constraints, Spapros selects genes that describe the overall variation in the scRNA-seq reference using a PCA-based selection procedure. To ensure high performance on cell type classification genes from a reference forest based selection are added. The pipeline takes into account prior knowledge like pre-selected genes, curated marker lists, and experimental constraints. **b**, Schematic of the transcriptome wide probe design pipeline. Genes for which not enough probes can be designed are filtered out, prior to gene set selection (first step in **a**). For the selected gene set technology specific ready-to-order probes are designed (final step in **a**) (created with BioRender.com). **c**, UMAP comparison of probe sets selected with Spapros for 50 and 150 genes and reference of 8000 highly variable genes for the Madissoon2019 human lung dataset. **d**, Dotplot of probes selected on the lung data set. Genes are ordered by the Spapros ranking system based on feature importance (see **Methods**). For each cell type the genes that are important for cell type classification based on the forest classification step are highlighted (Spapros marker). A minimum number of markers per cell type (DE or lit. gene) defined by the user is selected. For cell types not found in the dataset genes from a curated marker list are added. **e**, Difference of cell type classification confusion matrices between gene sets of Spapros and DE selections. AT1: Type I alveolar cell; AT2: Type II alveolar cell; DC1: Type 1 dendritic cell; DC2: Type 2 dendritic cell; NK: Natural killer cell; T CD4: CD4+ T cell; T CD8 Cyt: Cytotoxic CD8+ T cell.

As a first step, Spapros filters the full list of possible genes to exclude genes for which probes cannot be designed due to technology-specific technical constraints (Fig 2b). These constraints include the availability of sufficient unique possible probe sequences, as well as sequence properties like GC-content and melting temperature requirements. Binding locations of the final probes for a given gene cannot overlap. Thus, we generate non-overlapping probe sets with optimal thermodynamic and sequence properties with a graph-based search algorithm (see **Methods**). After ensuring that all remaining genes represent feasible probe candidates, Spapros selects genes that describe the overall variation in the scRNA-seq reference using a PCA-based selection procedure (see **Methods**) on a pre-selection of highly variable genes. To ensure cell types can be recovered using the probe set, Spapros uses the PCA-selected genes to predict cell type labels using a binary classification tree for each cell type (see **Methods**). The genes used in these trees represent candidate cell type marker probes, and the tree itself provides a combinatorial rule, describing how the cell types can be identified in the generated spatial transcriptomics data. To ensure that all cell types can be identified, Spapros compares the classification performance for each cell type to the performance of trees trained on DE genes, which are generated via a custom approach that optimizes for classifying similar cell identities (see **Methods**). If any discrepancy in performance is found with the DE trees (that represent the optimal performance target), Spapros iteratively adds DE genes to the list of possible probes to improve classification performance. Finally, genes are ranked based on their feature importance in classification trees to allow for a user-defined number of selected genes. To account for technical constraints of expression levels a smoothed multiplicative penalty kernel is applied to the scores of PCA and DE based selections (Fig. S4). Based on the final gene set and the non-overlapping probe sequences, our pipeline designs the final probe and detection sequences that need to be used for the experiment (see **Methods**).

While Spapros can select and design probe sets using only a reference scRNA-seq dataset and a list of cell types as input, users can also add prior knowledge and constraints to bias the algorithm toward user-defined probes. This allows users to add particular genes of interest (e.g., to test particular hypotheses or capture disease effects) and account for technological constraints (e.g., in situ sequencing has limitations on spot detection of highly expressed genes due to optical crowding). This prior knowledge can be incorporated in two ways: 1) as a pre-selection of probe genes, leading to other genes being combinatorially selected around them, and 2) as a marker list from which genes are added when respective cell types cannot adequately be classified (see **Methods**).

Overall, Spapros is a flexible, modular gene set selection and probe design tool that selects genes optimized for cell type recovery and cellular variation, while enabling users to customize the selection for any experimental design scenario.

### Spapros optimizes different probe set objectives simultaneously

We designed Spapros to optimize both cell type identification and recovery of variation beyond cell type annotations. However, by design, Spapros’ first priority is the identification of cell types, while more fine-grained variation is only captured if cell type identification is not affected. We therefore expect that for low numbers of genes Spapros mainly captures cell type level variation and gets more capacity for fine-grained variation with increasing numbers of genes.

Visually comparing the full gene set to Spapros selections on a UMAP embedding shows that cell type variation is strongly conserved using both 50 and 150 genes (Fig 2c, and Fig S8a). When comparing cell type classification characteristics between Spapros and DE genes for 50 genes we observe that similar cell types like DC1 and DC2 can be better distinguished by Spapros due to the combinatorial selection of e.g. CST3, FCER1A and IL1B (Fig 2d,e,S5,S8a). Spapros consistently outperforms the top performing classical feature selection methods DE and PCA on the prioritized objective of cell type identification, and optimizes variation recovery to the level of PCA-based selections when increasing the number of selected genes (Fig S8a). Thus, the Spapros probe set is optimized for the most relevant signals for any given number of selected genes.

### Spapros selection performs robustly across different datasets

When designing a targeted spatial experiment, data generators often have matching scRNA-seq data available from a matching sample. Yet, when this is not the case the question arises how similar the transcriptomic reference must be to the spatial sample. Using our evaluation metrics, we can address this question from a computational perspective. We assessed the cross dataset performance of probe sets selected on three different lung data sets (Fig S6a). Selections are not robust if the cell type classification and variation recovery performance shows high variance across data sets. To estimate if the variance across data sets is high or low we added selections on each individual donor sample as baseline comparisons.

We observe that the performance variance across data sets and across samples within each dataset are similar (Fig S6b). Thus, selections on one dataset show robust performance on other similar datasets. This is especially pronounced for cell type classification, indicating that there is no cell type identification performance drop for selections on an external dataset from the same tissue. As expected, the selection on the full dataset is beneficial for cell type classification compared to selections on individual donors. We also find this trend for variation recovery. Overall Spapros shows robust performance across different choices of the matched scRNA-seq reference for probe set selection.

### Spapros probe sets identify cell types and spatial variation within cell types in the adult human lung

When evaluated on scRNA-seq data, the Spapros pipeline shows high performance in cell type classification and reconstruction of variation beyond cell types. To show that these capabilities translate to spatial measurements we designed and performed a targeted spatial experiment using SCRINSHOT^13^ with a 64 gene probe panel generated by Spapros on healthy human lung samples using the Meyer2022^36^ scRNA-seq reference (see **Methods**). For each cell type Spapros provides a decision tree that includes the most important genes to robustly identify the given cell type. Leveraging these rules we detected all targeted cell types in an intralobar section (Fig 3a,b and S7; see **Methods**). Overall, the expression profiles of the cell type clusters matched the scRNA-seq reference clusters (Fig 3a) and the spatial distribution of cell types corresponded to known cellular structures in the lower airways and alveolar space (Fig 3b).

**Fig. 3:**
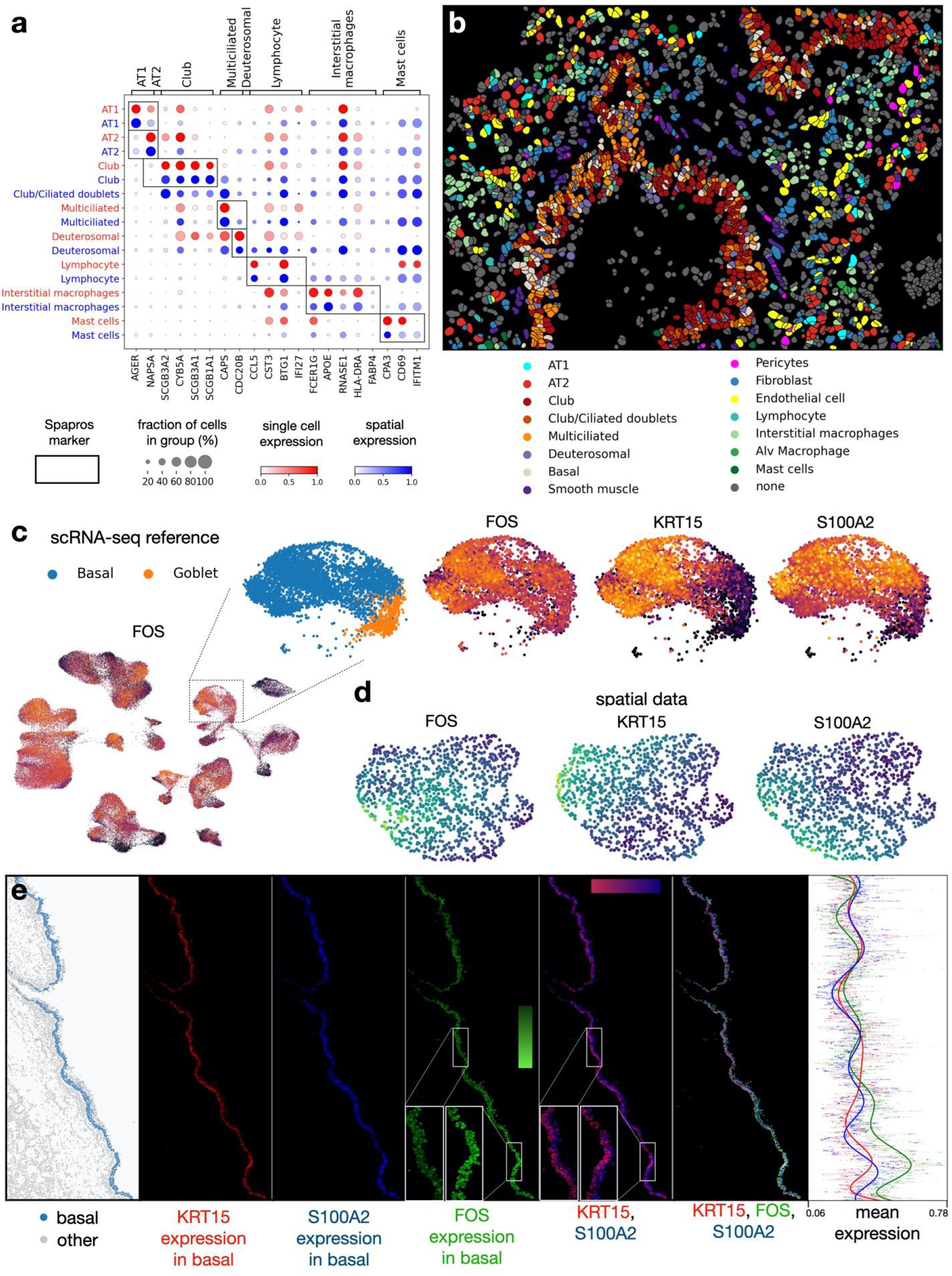
Spapros probe sets identify cell types and spatial variation within cell types. Spatial lung data measured with the SCRINSHOT technology for a probe set selected with Spapros. **a**, Mean expressions in spatial cell types of an intralobar lung sample (blue) and in cell types of the single cell reference (red). The shown genes are identified as the most important genes for cell type identification in the Spapros selection. **b**, Annotated cell types in the intralobar lung sample. **c-e**, Spatial distribution of two orthogonal variation axes within tracheal basal cells. **c**, FOS expression in the UMAP of the scRNA-seq reference data set, and expressions of FOS, KRT15, and S100A2 in the zoomed in basal and goblet subset. **d**, UMAP and **e** spatial distribution of FOS, KRT15, and S100A2 of the basal cells in a tracheal lung sample.

To capture variation beyond cell type annotations Spapros selects genes that exhibit gradients in multiple cell types as observed e.g. for FOS in our panel (Fig. 3c). Although FOS is typically associated with dissociation-induced cell stress^37^, we tested whether it also exhibits these gradients in non-dissociated tissue. Indeed, we observed such a gradient in airway epithelial cells. Here, FOS expression in tracheal basal cells displays intra-cell type spatial variation. While the basal cell markers KRT15 and S100A2 mark the inner (basal) and outer (suprabasal) epithelial layer respectively (Fig. 3c,d,e), FOS exhibits up- and down-regulated regions along the epithelium orthogonal to the interior-to-exterior epithelial variation of KRT15 and S100A2 (Fig 3d,e).

While further experiments are required to interpret this FOS signal, the signal itself indicates that probes selected by Spapros detect spatial variation of gene expression also beyond differences in cellular composition. Thus, Spapros enables identification of cell types of interest while also detecting spatially patterned intra-cell type variation in spatial measurements.

### Spapros outperforms curated gene sets and state-of-the-art methods

We assessed Spapros performance against 2 popular approaches for feature selection (DE and PCA), a curated gene list of airway cell type markers (see **Methods**), a published probe list for the human heart, and 8 recently proposed probe selection methods. The approaches were compared on two tissues: the heart and the lung, using publicly available scRNA-seq reference atlases and an untargeted spatial transcriptomics dataset (see **Methods**). Evaluation of all selected probe sets was performed on the basis of our evaluation suite. Spapros distinctly outperformed manually selected probe sets and curated marker lists in both heart^3^ and lung^36^ scenarios. Although manual selection of probe sets is commonly used^3–5,18^, Spapros outperforms these lists in both cell type identification and variation recovery (Fig. S8b,c). Notably, the probe set for the developmental heart was generated for an in situ sequencing (ISS) experiment on the basis of both an scRNA-seq reference and an untargeted *spatial transcriptomics*^*7*^ assay. Adapting Spapros to leverage both of these data sources (see **Methods**), we find that our optimized pipeline outperforms the curated probe sets in all evaluated metrics.

Recently published computational methods for probe set selection were compared in a large-scale benchmark on both heart and lung datasets, while the heart data has more fine-grained cell type annotations compared to the lung data. All methods were run to generate both a small (50 genes) and large (150 genes) probe set (Fig S8a). Consistent with the method designs, the results show that methods can be grouped into categories with differing goals: general variation recovery (selfE, scmer, PCA, triku), cell type identification or selection of cell type specific markers (smash, nsforest, asfs, scgenefit, cosg, DE), and both of these objectives (Spapros).

We find that across tasks Spapros performs best in cell type classification. This is especially pronounced for low numbers of genes (50) where Spapros differentiates more strongly from other methods. For a target probe set of 150 genes, the methods that optimize for cell type identification improve classification performance, while only nsforest consistently performs as well as Spapros across tasks. The variation recovery methods (selfE, scmer, PCA) perform best at capturing the general transcriptomic variation in the data with a reduced probe set. While Spapros scores 5% worse than the top performer for 50 probes, it performs similarly for a target of 150 probes despite concurrently optimizing for cell type identification. In contrast, other cell type identification methods do not optimize for variation recovery at 150 genes, and other variation recovery methods do not reach the cell type classification performance of Spapros. Furthermore, none of the algorithms we compare with our method is able to account for technical constraints.

Overall, we find that Spapros is the only method that performs well on both cell type identification and variation recovery. Thus, Spapros provides a multi-purpose end-to-end probe set selection pipeline that can use both scRNA-seq and untargeted spatial transcriptomics data to optimize gene set selection and probe design.

## Discussion

We present Spapros, a probe set selection and design pipeline for experimental design of targeted spatial transcriptomics experiments. Spapros optimizes probe set selection in a combinatorial fashion through optimizing both design and selection of genes simultaneously while taking into account prior knowledge and technical constraints. With these features our method is the first one that enables end-to-end probe set selection. It is also the first method that optimizes simultaneously for both identification of cell types and recovery of transcriptomic variation beyond cell types.

Due to a reduced number of measured features and processing challenges, annotation of cell types is typically more challenging in targeted ST data compared to scRNA-seq data. Thus, currently available ST studies generally focus on cell type identification in space and most available probe selection methods focus on that objective. Spapros facilitates cell type identification not only by optimizing probe sets for this purpose, but also by providing rules for cell type annotation based on the decision trees used for probe selection. In this study, we used these annotation rules to identify lung cell types in a SCRINSHOT experiment. Furthermore, Spapros also highlights which probes are candidates for spatially variable patterns across cell identities by scoring probes on variation recovery and their importance for cell type separation. These hints led us to the investigation of FOS which shows spatial intra-cell type patterns along the airway epithelium. It is valuable to capture intra-cell type variation since meaningful spatial signals go beyond differing cell type compositions across locations as they can indicate local tissue niches, viral spread, inflammation, and more. However, including genes like FOS reduces the separation of cell types when clustering cells. Thus, such genes introduce an additional challenge for downstream analysis and should be excluded for cell type clustering. Spapros’ rule-based annotation output and labeling of gene characteristics enable a quick reference-based spatial mapping of cell types and identification of new spatial patterns of cell state continuums. As spatial transcriptomics protocols and analysis methods continue to improve, these spatial state continuums will become increasingly important.

A central assumption in the current implementation of Spapros, and probe set selection in general, is that the transcriptomic signal measured in scRNA-seq is representative of the signal measured in targeted spatial transcriptomics. As shown in our lung study we are able to find the cell types of interest based on reference signals with high recall and we can find matching orthogonal cellular variation within cell types. However, the equality assumption between spatial and scRNA-seq data is currently not strictly met and we find discrepancies between the modalities. Gene distributions are notably different and some genes are uncorrelated to the signal expected from scRNA-seq, possibly due to non-functional probes or individual tissue section anomalies. There are multiple reasons that lead to discrepancies between the modalities: Different RNA measurement techniques lead to the exclusion of reads from highly similar paralogues that are mapped to multiple regions of the genome in droplet based scRNA-seq; Different sample processing between scRNA-seq and targeted ST lead to the enrichment or exclusion of certain cell types and states.; and technology-specific effects like optical crowding can result in imprecise measurements. Quantifying these differences and constructing additional robustness constraints are important future directions for probe set selection and reference based down stream analysis. These robustness constraints also depend on the processing of targeted ST data for which there are currently no best practices. Normalization, further pre-processing, and cell segmentation including the disentanglement of overlapping cells will affect how well these modalities match. To design ideal robustness constraints future studies need to investigate these effects as well as tissue parameters like organ type, cell density, and tissue quality.

Further improvements to probe set robustness can be derived from integrated reference atlases. As more scRNA-seq datasets are becoming available, these datasets are being integrated into comprehensive reference atlases^38^ that contain consensus signatures of rare cell types and subtle state differences learned across studies. Yet, the large cell number also poses additional methodological challenges such as batch effects and unbalanced label distributions. Spapros tackles unbalanced label distributions by balanced sampling strategies over cell type labels to robustly capture rare cells. Yet, batch effects are still a challenge for Spapros and other probe selection methods. Especially selection methods that optimize for variation recovery will also select genes aligned to batch effect variation. While we showed that Spapros can select probe sets and project these across datasets, we did find that also dataset-specific variation (potentially due to batch effects) was captured. The optimization for biological variation disentangled from technical variation is therefore an interesting direction for future work. Further extensions to Spapros include experimental design for other modalities like spatial proteomic measurements (e.g. Codex^39^).

With Spapros we introduced new concepts for optimally selecting probe sets in targeted spatial transcriptomics: our approach combines gene panel selection and probe design to enable combinatorial selection, and optimizes simultaneously for the dual objective of cell type identification and recovery of transcriptomic variation. Spapros will thus enable optimal experimental design while guiding downstream analysis. Additionally our evaluation suite sets a reproducible and robust standard for quality assessment of spatial probe sets and can be readily extended towards additional metrics. Spapros is available as a Python package enabling easy and flexible probe set selection, evaluation of probe sets, and probe design. With Spapros we aim to enable users to maximize their success in future exploratory spatial studies to find novel spatial cellular variation.

## Methods

### Probe set evaluation metrics

We assessed the selection method performance or probeset quality via multiple metrics of the categories cell type identification, variation recovery, gene redundancy, fulfillment of the technical constraints, and computation time. For the calculation of the metrics a reference dataset (typically scRNA-seq) is used. The reference is reduced to a pre-selection of highly variable genes (8000 in our use cases; scanpy’s highly_variable_genes function with cell_ranger flavor option). In the following the metrics of each category are described.

#### Variation recovery

For the clustering similarity metrics Leiden clusterings for different numbers of clusters *n*_*c*_ are calculated via binary search over Leiden resolutions. This way sets of clusterings for *n*_*c*_ ∈{7,8,…,60} are produced for the 8000 reference genes and the gene set that is evaluated. For each *n*_*c*_ the similarity of the clusterings between reference and gene set are measured via normalized mutual information:

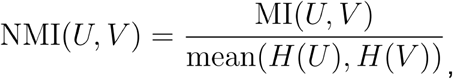

with *U*: set of sets of cells in each cluster 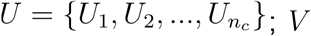; *V*: like *U* for the second clustering; *MI* : mutual information which is defined as

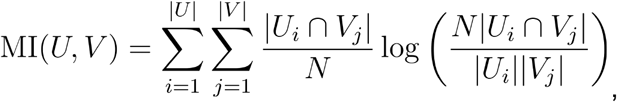

With *N* : number of cells; note that |*U*| = |*V*| = *n*_*c*_ in our comparisons; *H(U)*: entropy of *U* which is given by

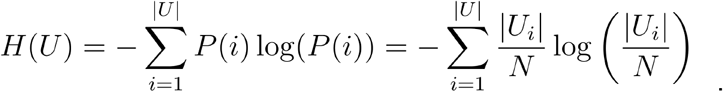

The final clustering similarity metrics are given by

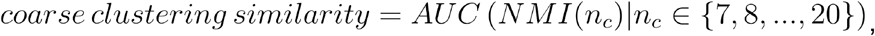

and

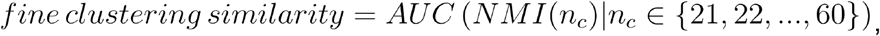

with 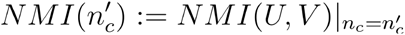, and *AUC*(*f*(*x*)|*x* ∈ I) : the area under the curve of *f(x)* over the interval. Due to the nature of the Leiden algorithm sometimes certain for a given dataset can not be found. Missing values due to that are imputed by linear interpolation between *NMI*(*n*_*c*_−1) and *NMI*(*n*_*c*_+1) or the closest existing values in case of multiple missing data points.

For the n*eighborhood similarity* metric knn graphs are obtained for the 8000 reference genes and the gene set that is evaluated. Knn graphs are calculated for *k* ∈ {5,10,20,30,50} on the PCA space of the gene expression. The n*eighborhood similarity* is given as

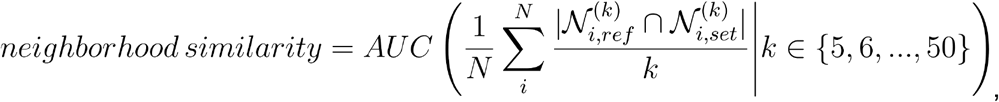

with *N* : number of cells, and 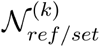 : set of neighbors of cell *i* in the knn graph of the reference (ref) or gene set (set) for a given *k*.

#### Cell type identification

To assess how well cell types can be recovered with a gene subset we train gradient boosted forests^40^ for multi-class cell type classification and measure the test set performance. To achieve a robust performance readout 5-fold cross validation is performed with 5 different seeds, summing up to *N*_*m=25*_ models per evaluated gene set. The test set classification confusion matrix of each model is obtained and normalized by the ground truth cell type count. Based on the normalized confusion matrix *CM*_*m*_ per model *m* the summary metrics are given as

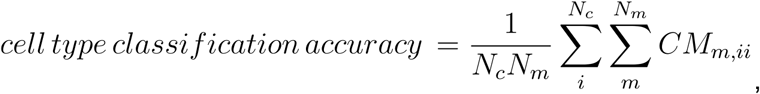

and

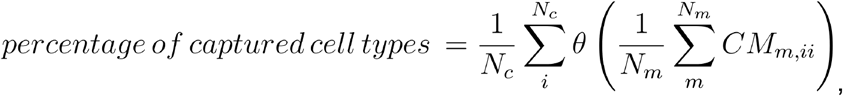

with the linearly smoothed step function *θ*(*x*) from 0.75 to 0.85 (i.e. *θ*(*x* ≤ 0.75) = 0 and *θ*(*x* ≥ 0.85) = 1), and the number of cell types *N*_*c*_.

The metrics *marker correlation* and *cell type balanced marker correlation* measure how well marker signals of a literature derived marker list are captured with the selected gene set. Based on the maximal pearson correlation 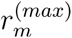 with the gene set for each marker *m* the summary metrics are given by:

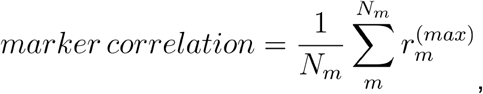

and

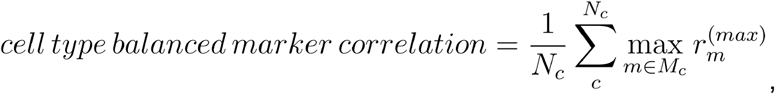

with the number of markers *N*_*m*_, the number of cell types *N*_*c*_, and the set of markers *M*_*c*_ for each cell type *c*.

#### Gene redundancy

Based on pearson correlations *r*_*ij*_ of gene pairs (*i,j*) we asses the redundancy in a gene set with the overall

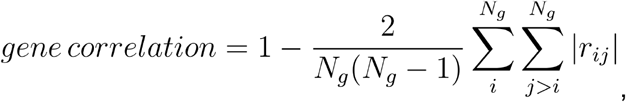

and the

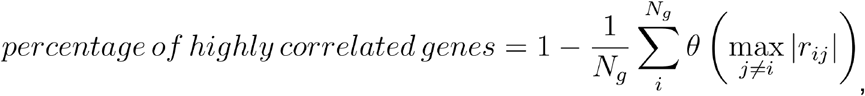

with the number of genes *N*_*g*_ in the gene set and the linearly smoothed step function *θ* (see above).

#### Expression constraint violation

We penalize a gene for too low expression if it is expressed below a lower expression threshold in at least 90% of cells where the gene is expressed > 0, i.e. the 0.9 expression quantile. Similarly we penalize too high gene expression if the 0.99 expression quantile over all cells is above an upper threshold. Expression thresholds were obtained from expert experience on too lowly and too highly expressed reference genes (e.g. MALAT1). Based on cpm log-normalized data the thresholds were set to 2.3 and 5. Since our data was scran normalized thresholds were transferred by mapping mean expressions to 1.78 and 4.5. These values are technology specific and can only be roughly estimated. To not set strict thresholds, smoothed penalty functions *P*(*q*) over the gene’s quantile *q* with a Gaussian decay below (low expression) and above (high expression) the thresholds were introduced. To assess how strongly a gene set violates the expression constraints we compute the means over *P*^*low*^(*q*) and *P*^*high*^(*q*) as the *low* and *high expression constraint violation* metrics respectively.

### Classical feature selection methods

The evaluated classical feature selection methods include differentially expressed (DE) genes selection, a PCA-based selection, highly variable genes (HVG) selection, a sparse-PCA (SPCA) based selection and selection of highly expressed genes. DE genes were scored with t-tests using scanpy’s ^41^ rank_genes_groups function. The highest scored genes per cell type were selected till the set encompassed *n* genes. For the PCA-based selection genes were scored according the sum over the loadings of the first 20 PCs, and the top *n* genes were selected. HVG selection was performed with scanpy’s highly_variable_genes function and the cell_ranger flavor option. For the SPCA-based selection scikit-learn’s sparsePCA class was used. With the *α* argument of sparsePCA the sparsity of the loadings matrix can be controlled. To select as set of *n* genes a binary search over *α*’s was conducted to find a setting where only *n* genes have loadings > 0. For the selection of highly expressed genes we scored genes based on the mean expression.

### Spapros selection pipeline

Spapros enables an end-to-end probe set selection which include probe design constraint based gene filtering, gene panel selection, and the probe design for the selected gene panel. Therefore Spapros encompasses a probe design pipeline and a gene set selection pipeline. We discuss them separately in the following.

#### Gene panel selection

Spapros uses a log-normalized count matrix of all genes or a pre-selection of highly variable genes and cell type annotations as input. The first step consists of a PCA-based (see classical feature selection methods) prior selection (100 genes per default). This prior selection biases the next steps to use genes that capture a high degree of variation in the dataset. Next we train decision trees on the prior selected genes for binary cell type classification for each cell type (i.e. cell type of interest vs all other cell types). These trees are highly regularized by setting the max_depth to 3. Thus, robust and interpretable rules are learned for the classification of each cell type. For each tree optimization a training set of 1000 cells per cell type are sampled (oversampling if cell count is too low), and performance is assessed on a test set sample of 3000 cells per cell type. The uniform sampling ensures cell type balanced training. Additionally we use class proportion weights for the binary classification of the two classes “cell type of interest” vs “other”. Per default we train 50 trees per cell type each with a different training set and choose the best tree based on the F1-score. To increase classification performance on cell types that are hard to distinguish from the target cell type we train additional secondary trees on a subset of cell types. This subset is identified based on the specificities *s*(*c*) = *TN*_*c*_/*N*_*c*_ of the primary tree of each cell type *c* in class “other”, with *TN*_*c*_ : number of true negative cells of cell type *c*, and *Nc* : number of cells from cell type *c* in test set. Cell types are considered for the secondary tree training if their specificity is either below 0.9 or one standard deviation below the mean of all specificities, but at least 0.02 below the mean. Spapros has a hyperparameter to set the number of further secondary trees that are trained based on previous secondary trees in the same manner. The default is 3, i.e. 1 primary and 2 secondary trees. The trees trained on PCA selected genes often can be improved since important genes in the pool are missing. Therefore we also train trees on DE genes. Trees are trained in the same manner except that additionally an iterative adding of genes from specific DE tests is performed: After tree training cell types that are hard to distinguish are identified via specificities (best over primary and secondary trees) the same way as cell types were selected for secondary trees. Then a DE test is performed between the cell type of interest and the identified cell types. The top 2 of those DE genes per cell type that needs optimisation is added to the DE pool and tree training is repeated till an early stopping criterion is reached or up to 12 times. By comparing the performance of trees on PCA genes and trees on DE genes we identify those cell types that are better distinguished with DE genes and iteratively add missing genes from DE trees with highest feature importance to the PCA pool and retrain trees till the same performance is reached. The genes that occur in the final trees are ranked by feature importance and build the gene panel.

Spapros incorporates prior knowledge as a pre selection of genes that is added to PCA and DE pools, so genes are selected around them. Further a list of marker genes can be provided. After tree based selection the pipeline checks if at least a user defined number of markers per cell type are captured (correlation > 0.5; see marker correlation metrics). Experimental expression constraints are incorporated by multiplying PCA and DE scores (see classical feature selection methods) with a penalty kernel (see expression constraint violation metrics).

#### Probe design pipeline

To generate a genome wide probe design filter and to design probes for a set of given genes, we developed a custom probe design pipeline. Currently the pipeline supports padlock probe design for SCRINSHOT, however many steps are shared between different technologies. The pipeline has four major steps: 1) Create probe and background databases, 2) Filter probes by sequence property and binding specificity, 3) Rank sets of non-overlapping probes for each gene and 4) Generate ready-to-order padlock probe sequence.

To create the probe and background databases, the user has to define the *species, genome assembly, annotation source* and *annotation release* for the reference genome. As annotation source, the user can choose between *NCBI* and *Ensembl* annotation. The genome sequence (fasta format) and gene annotation (gtf format) files are automatically downloaded via the respective FTP server. If the annotation source is NCBI, the chromosome names are automatically mapped from *GenBank* to *RefSeq* annotation in order to be used by bedTools. Alternatively, the user can provide a custom genome sequence (fasta format) and custom gene annotation (gtf format). To create the probe database, the user has to define the *probe length*, which can be given as a range, and provide a list of genes for which probes should be created. The *gene list* should be a text file where each line corresponds to a gene identifier and the identifier has to follow the annotation source standards. If the user does not provide a list of genes, the pipeline will create probes for all genes in the gene annotation file. The probe database contains all possible probe target sequences for each gene in the gene list. The target sequences are created from a sliding window over the DNA sequence of the genes coding strand. Probe target sequences are created from the transcript sequences of all existing isoforms for a gene. The background database contains the full transcriptome of the provided annotation, i.e. all transcript isoforms for all annotated genes, including protein-coding and non-coding RNAs. To speed-up downstream processing steps (e.g. BlastN search), redundancies within the transcriptome, e.g. isoforms using several common exons, are minimized. The minimization of redundancies is commonly achieved through multiple sequence alignment. To avoid the time-consuming step of a multiple sequence alignment over all transcripts, a reduced transcriptome is created from the gene annotation itself. The reduced transcriptome consists of all annotated exons and all possible exon junctions defined by the different transcript isoforms. An exon junction region is defined as the region *probe length + 5 bp* upstream of the first exon and *probe length + 5 bp* downstream of the second exon. The exon junction region is larger than the probe length (+ 5 bp) to allow bulges in alignment calculations (e.g. BlastN). The resulting transcriptome annotation contains each exon and all possible exon junctions only once per gene, i.e. when multiple transcripts use the same exon, the region is only reported once. The transcriptome annotation is saved in *bed12* format that allows split annotations, which are needed to get sequences for exon junctions, i.e. the intron sequence has to be skipped. The *fasta* transcriptome sequence is retrieved from the *bed12* file using bedTools getfasta.

After creating both databases, each target sequence in the probe database is filtered by certain sequence properties, i.e. by undefined nucleotides in their sequence (masked with ‘N’) as well as *GC content* and *melting temperature* for user-defined ranges. Specifically for padlock probes a ligation site needs to be set that generates two probe arms. The pipeline searches for a ligation site that generates two arms with a *maximal melting temperature difference* of 2°C and filters probes if such ligation is not found. The probes passing all sequence property filters are aggregated over each gene, i.e. all probes of one gene having the exact same sequence are merged into one entry, saving the information of exon and transcript-of-origin as well as start and end position of the respective probes. After applying the sequence property filter, a specificity filter is applied, which removes probes that potentially bind to other similar transcripts, so called off-targets, excluding matches to the probes gene. Off-targets are identified through an alignment search with BlastN. BlastN alignment finds regions of local similarity between query and target sequences, where the query sequence are sequences of the probe database and the target sequence are sequences of the background database. Before running BlastN, a pre-filtering is applied to remove all duplicated sequences (exact matches) within the probe database. The user should define the *word size* for the BlastN search, which defines the length of the BlastN seed region. BlastN returns query-target pairs, where both sequences match with a certain *coverage* and *similarity*, for which the user has to define thresholds. Probes are then filtered based on the percent identity, the alignment length (coverage) and optionally the coverage of the region around the ligation site of the probe by the target sequence, where the user has to define the ligation region.

Next, the pipeline searches for the best set of a user-defined *number of non-overlapping probes*. Sets are scored by the highest distance to optimal GC and melting temperature of the probes in the set. Based on genomic locations an overlap graph for the probes of a given gene is generated. The pipeline iterates through non-overlapping sets that are given by the cliques of the complement graph, measures their score, and ranks sets. If there are many probe candidates this search can take long. Therefore the search is sped up via a candidate filter. For that filter the best probe is selected, then the next best that doesn’t overlap with the first, and so on, till the number of non-overlapping probes is reached. This is repeated 10000 times starting with the 2., 3.,… best probe. For the clique based search all probes that score worse than the worst probe of the best set of the heuristic search are filtered out.

Finally padlock probes and detection oligos are designed for the best non-overlapping sets of each gene. For this adding the backbone, pruning the oligo sequence for optimal melting temperature, exchanging Thymines to Uracils in the detection oligo, and placing the fluorescent dye at the side with the closest Uracil as described in^13^ was automated.

### Datasets for probe set selection and evaluation

Our experiments and analyses comprise three human lung scRNA-seq datasets *Madissoon2020^42^, Krasnow2021*^*43*^, *Meyer2022*^*36*^, a scRNA-seq dataset and an untargeted spatial transcriptomics dataset of the developing human heart *Asp2019 (sc/ST)*^*3*^, and a sc/snRNAseq adult human heart dataset *Litvinukova2020*^*44*^. The datasets are all publicly available. Cell type annotations were obtained from the original publications. For fair comparisons the annotations were filtered or pooled to coarse annotations in those analyses where necessary (Table A1). All single cell and single nucleus datasets were preprocessed in the same way: raw counts were normalized with scran^45^ using Leiden clusterings^46^ with resolution 0.5 on a temporary log normalization to 10^6^ counts per cell. The logarithm of the scran normalized data plus one pseudocount was taken. Features were reduced to the top 8000 highly variable genes selected with scanpy’s highly_variable_genes function (flavor: cell_ranger). A detailed summary which dataset and annotation was used in each analysis is given in Table A1. For some datasets we only used a subset of cells due to different reasons: For Meyer2022 we only used the single cells and not nuclei since we used it for our SCRINSHOT experiment and assume that single cell expressions are closer to the observation in a targeted ST experiment compared to single nuclei. Only the developmental stage at 6 weeks from Asp2019 ST was used since the selected ISS panel in their study was also selected on that subset. The heart atlas Litvinukova2020 was reduced to the single nucleus observations and maximal 2k cells per cell type to reduce computation time of our evaluations (56 cell types, including 58966 cells).

### SCRINSHOT experiment

#### Samples and histology

Samples were obtained from deceased transplant organ donors by the Cambridge Biorepository for Translational Medicine (CBTM) with informed consent from the donor families and approval from the NRES Committee of East of England – Cambridge South (15/EE/0152). Lung biopsies (~2cm^3^) were fresh-frozen in OCT (Leica Surgipath, FSC22), and shipped to Stockholm University on dry ice. Quality control was carried out by evaluating histopathological condition (sections stained with hematoxylin and eosin were analyzed by the pathologist) and RIN value analysis. Healthy samples with RIN values above 4 were selected for SCRINSHOT. Sections of lung tissues were cut at 10 μm thickness and placed on poly-lysine slides (Thermo, J2800AMNZ), then stored frozen at −80°C for further use.

#### Probe design

At the time when the probe set for the SCRINSHOT experiment was selected with the Spapros gene panel selection the probe design pipeline was not finished, therefore probes were designed manually. A detailed description of the padlock probe design is provided in previous publications ^13,47^. The sequences for probes (38-45 nt length) were selected using PrimerQuest online tool (Integrated DNA Technologies: IDT) for the targeted mRNA of 64 genes in the gene selection list. These sequences were then interrogated against targeted-organism genome and transcriptome, with Blastn tool (NLM) to guarantee their specificity. Two to four specific sequences per gene were selected for further padlock design. An extra sequence was selected to create a unique barcode for the detection probe, which was re-used for each padlock of the same gene with several T nucleotides replaced with U, as described previously ^13,47^. All detection probes were then interrogated against all padlock probes using Blastn tool in order to ensure no overlapping sequences and avoid unspecific detection probe binding. An overlap of nine or more nucleotides was avoided by modification of the detection barcode by replacing 1-2 nt. One RCA primer sequence was used for all padlock probes taking into account the preceding gene expression level pre-selection. Sequences for padlock and fluorophore-labeled detection probes are provided in Table A2. Both types of probes (248 padlock and 64 detection probes) were ordered from IDT.

#### SCRINSHOT procedure

SCRINSHOT procedure was followed exactly as described previously ^13,47^ with extra stringent detection probe incubation (30 °C) in 30% formamide, and increased (20%) concentration of formamide of washing buffer in the following step in order to avoid unspecific binding of detection probes. After a trial experiment, SCGB1A1 and SCGB3A1 probe concentration was reduced to one padlock per gene to avoid dot crowding. Probes were applied in sets of five per hybridization cycle, a total of 13 cycles to detect all 64 genes in each sample (Table A3). After each cycle the whole slide was imaged as a Z-stack with 11 steps of 0.8 μm (to cover the whole 10 μm thickness) using a widefield microscope (Zeiss Axio Observer Z.2, Carl Zeiss Microscopy GmbH, with a Colibri led light source, equipped with a Zeiss AxioCam 506 Mono digital camera and an automated stage) at 20x magnification. Maximum intensity orthogonal projection was then used for further analysis as described previously 13,47. One sample from an area corresponding to alveolar parenchyma collected from the upper part of the left lobe (Location 5, Luecken et al, 2022^48^) with a substantial amount of signal from most of the probes was selected for gene pre-selection evaluation.

#### SCRINSHOT image analysis

Image alignment using DAPI channel, followed by manual nuclei segmentation for intralobar region, and automated nuclei segmentation for tracheal region. For the automated segmentation a MaskRCNN convolutional deep neural network model was used as part of the NucleAIzer pipeline. The final model is trained so that the annotated image set was augmented with artificially created ones ^49^. The training set contained 50.000 single nuclei manually annotated by experts, on 40x magnification microscopy images. The trained network was integrated into the BIAS (Biological Image Analysis Software) ^49,50^ and is available in: http://single-cell-technologies.com/download/. Automated dot detection using CellProfiler was performed as described previously ^13,47^. All detected dots were assigned to each cell ROI in Fiji (https://github.com/AlexSount/SCRINSHOT/blob/master/automated_stitching_dot_counting_v1_19genes.ijm). The resulting dataset containing dots per ROI was used for further analysis.

#### SCRINSHOT analysis

Cells with less than 10 counts were filtered out. Counts were normalized by segmentation area and then logarithmized. The cells were clustered with the Leiden algorithm and cell types were annotated by comparing expression profiles of Spapros markers for each cell type with the Meyer2022 scRNA-seq reference. Inclusion of some genes affected the clustering in a worse separation of cell types. Those genes were therefore left out for the clustering (Table A4). These genes include broadly expressed genes with intra-cell type variation like FOS. They were identified based on the PCA-scores in the Spapros selection and manual inspection of mean expressions over cell type clusters.

For the trachea sample, only genes that were relevant for identification of basal cells were included in the clustering (Table A4). We searched for genes with orthogonal intra-cell type variation to KRT15 and S100A2 based on low prediction scores of a linear regression on KRT15 and S100A2 and high abundance in basal cells. FOS turned out as a strong candidate in comparison of all genes. For the smoothed spatial expression profile of KRT15, S100A2 and FOS along the epithelium scikit-learn’s B-spline fit with 10 knots, degree 10 and l2 regularization with *α* = 10^−3^ was used.

### Selection method benchmark

#### Curated marker list and ISS panel

The curated lung marker list (Table A5) was provided by lung experts (see acknowledgements) and is a collection of airway wall markers from various publications. We reduced the number of genes in the marker list to 155 by only allowing up to 10 genes per cell type and from those the ones that occur in the 8000 highly variable genes of the Meyer2022 dataset (see Datasets for probe set selection and evaluation).

For our comparison with an ISS panel we took the original gene set from Asp2019^3^ which contains 69 genes that were selected based on an scRNA-seq and an untargeted spatial transcriptomics data set. To generate a comparable selection with Spapros we selected 34 genes on the untargeted dataset and used these as prior knowledge selection (see Spapros selection pipeline, gene panel selection) for a selection of 69 genes on the scRNA-seq data set.

#### Method benchmark

Methods were benchmarked on the data sets Madissoon2019 and Litvinukova2020 (see datasets for selection and evaluation and Table A1)

As recommended for the method scmer, subsampling the dataset is necessary to run the method in a reasonable time. We followed their recommendations of sub sampling to 10000 cells. The method selfE takes even longer, therefore we applied the same sub sampling scheme as for scmer. If a selection took longer than two days it was interrupted and not added (selfE and asfs for 150 genes).

#### External selection methods

We compared Spapros with 8 other methods dedicated to gene selection. These methods are described in the following.

*NS-Forest*^*23*^ is a marker gene selection algorithm based on random forest importance scores combined with a binary expression scoring approach to select markers that are specifically up-regulated in the cell type of interest but not in other cell types. NS-Forest is available as a repository of python functions.

*SMaSH*^*21*^ is a general computational framework for extracting marker genes. Different base classification models can be used: three different forest based ensemble learners and a neural network. Gini importance and Shapley values are used for scoring genes for the forest models and the neural network respectively. As the authors describe that the XGBoost base model performs consistently excellent in terms of yielding low marker gene classification rates we chose this configuration for our comparisons. SMaSH is available as a python package on PyPI.

*scGeneFit*^*20*^ selects gene markers that jointly optimize cell label recovery using label-aware compressive classification methods. The method finds a projection to the lowest-dimensional subspace for which samples with different labels remain farther apart than samples with the same label, while the subspace dimensions are individual genes. The optimization is formulated as a linear program. The method not only finds marker genes that are specifically expressed in single cell types but also genes that reflect the hierarchical structure of cell types. scGeneFit is available as a python package.

The *ASFS* (or *ActiveSVM*)^30^ selection procedure generates minimal gene sets from single-cell data by employing a support vector machine classifier. The method iteratively adds more genes by identifying cells that were misclassified. ASFS is available as a python package on PyPI.

*COSG*^*25*^ is a cosine-similarity based method for marker gene selection. It is fast and scalable and particularly designed for selection of marker genes on large datasets. COSG is available as python and R packages.

*SelfE*^*31*^ aims to select a subset of genes that is optimized for prediction of all remaining genes as linear combinations. The gene subset is constructed iteratively and each step the gene that minimizes the l2 error over genes is added. SelfE is available as an R package.

*SCMER*^*32*^ selects a set of genes that reconstructs a pairwise similarity matrix between cells and therefore preserves the manifold of the scRNA-seq data. To find that sparse set of features a binary search on the l1 regularization parameter is performed. Similar to SelfE this method optimizes for general variation opposed to the previously described marker gene- and cell type classification-focused methods. The method is available as a python package. *Triku*^*34*^ selects genes that are locally overexpressed in groups of neighboring cells which aims to recover cell populations and general variation in the scRNA-seq data. The method is available as a python package.

## Supporting information

Supplemental Image Files

## Acknowledgements

We are grateful to all members of the Theis and Ertürk laboratories, as well as the discovAIR consortium for frequent discussions of the project. We thank Elo Madissoon and Kerstin Meyer for provision and discussion of the scRNA-seq lung reference datasets. We thank Pascal Barbry for the provision of the airway marker list. We thank Jonas Theelke for testing the probe design pipeline. We thank Xesus Abalo for helping with tissue sectioning and tissue quality control. We thank Wim Timens for histopathological tissue evaluation.

## Author contributions

MDL conceived the project. LBK, MDL developed the Spapros method and metrics. LBK conducted the computational experiments. MDL, FJT, CS supervised the project. AS and CS selected the parameters for gene and probe selection. LBdAeS, IIM, LBK set up the probe design pipeline. AS and ABF revised the probe design pipeline. KTM collected the tissue samples for SCRINSHOT. ABF prepared the samples for SCRINSHOT and conducted the SCRINSHOT experiment. TB, FK conducted the automated nuclei segmentation of the tracheal sample. LBK, ABF analysed the SCRINSHOT data. LBK, LS, LH contributed to the Spapros package. LH wrote the Nextflow evaluation pipeline. LS implemented the external selection methods. LBK, MDL, ABF, LBdAeS, FJT wrote the manuscript. All authors reviewed the manuscript.

## Data and code availability statement

The SCRINSHOT expression data is attached as an excel file. All sc/snRNA-seq and untargeted spatial transcriptomics datasets are publicly available^3,36,42−44^. The Spapros package and the probe design pipeline are publicly available at https://github.com/theislab/spapros and https://github.com/HelmholtzAI-Consultants-Munich/oligo-designer-toolsuite respectively. The end-to-end selection which combines panel selection and probe design is described in tutorials of the Spapros package. The Nextflow evaluation pipeline is available at https://github.com/theislab/spapros-pipeline. Code to reproduce the analyses will be available at https://github.com/theislab/spapros_reproducibility.

## Supplementary

### Tables

A1 Cell type annotations for all analyses

A2 SCRINSHOT probe sequences

A3 SCRINSHOT measurement cycles

A4 SCRINSHOT expression data

A5 Curated marker list of airway wall cell types

### Files

F1 SCRINSHOT cell segmentations intralobar region

F2 SCRINSHOT cell segmentations tracheal region

### Figures

**Fig. S1:**
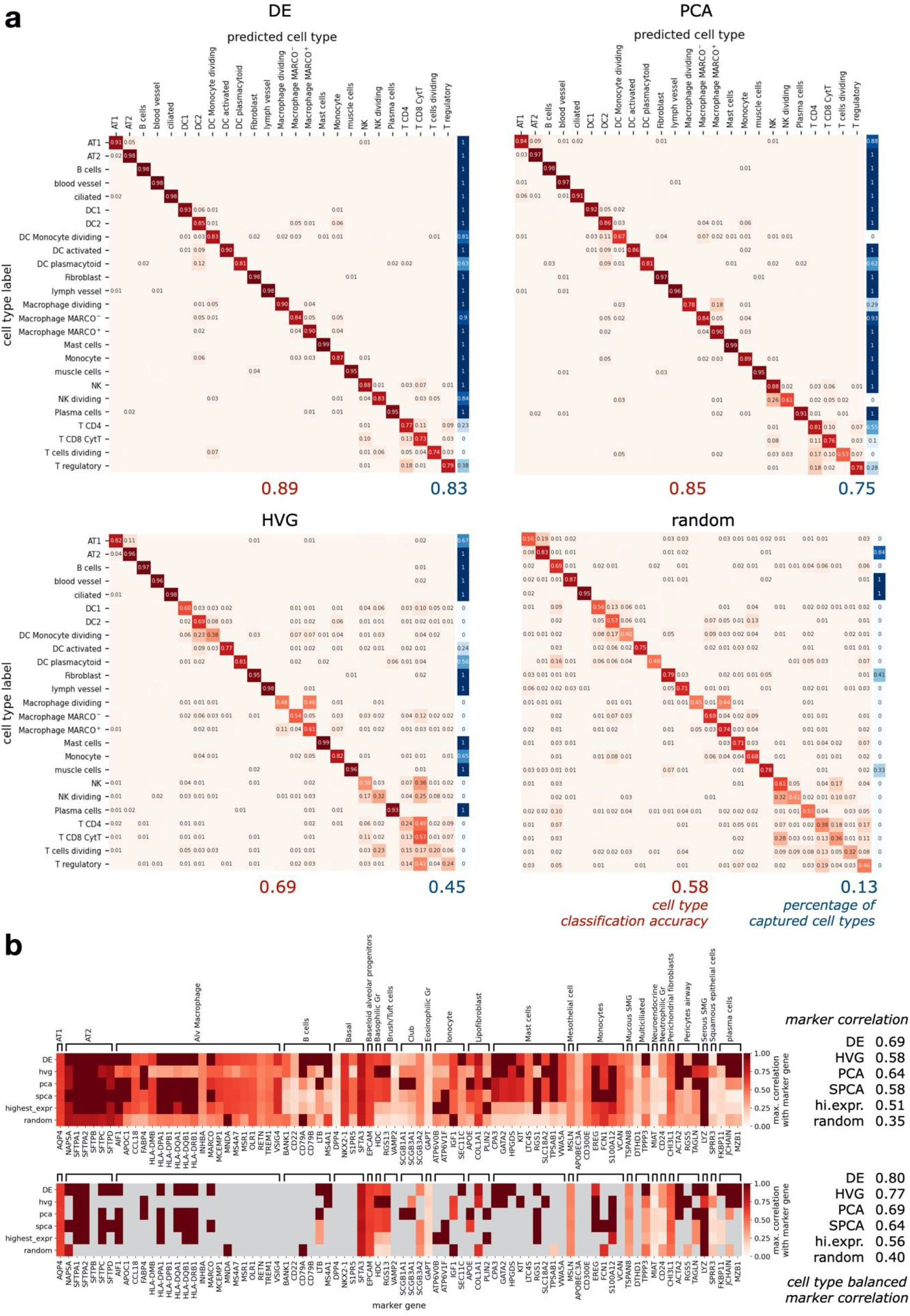
Spapros evaluations show cell type specific classification performance. Evaluations on the Madissoon2019 dataset. **a**, Normalized cell type classification confusion matrices (red color scale) for gene sets of 150 genes selected with DE, PCA, HVG, and random selection, and linearly smoothed step function of the diagonal elements at 0.8 (blue color scale). The summary metrics *cell type classification accuracy* and *percentage of captured cell types* are the means of the diagonal and the thresholded values respectively. **b**, Maximal Pearson correlation of marker genes from a curated marker list and gene sets selected with DE, HVG, PCA, SPCA, as well as highest expressed and randomly selected genes. In the bottom heatmap values below the maximum correlation of each cell type are masked (gray). The summary metrics *marker correlation* and *cell type balanced marker correlation* are the row means of all genes (top heatmap) and per cell type (bottom heatmap) respectively.

**Fig. S2:**
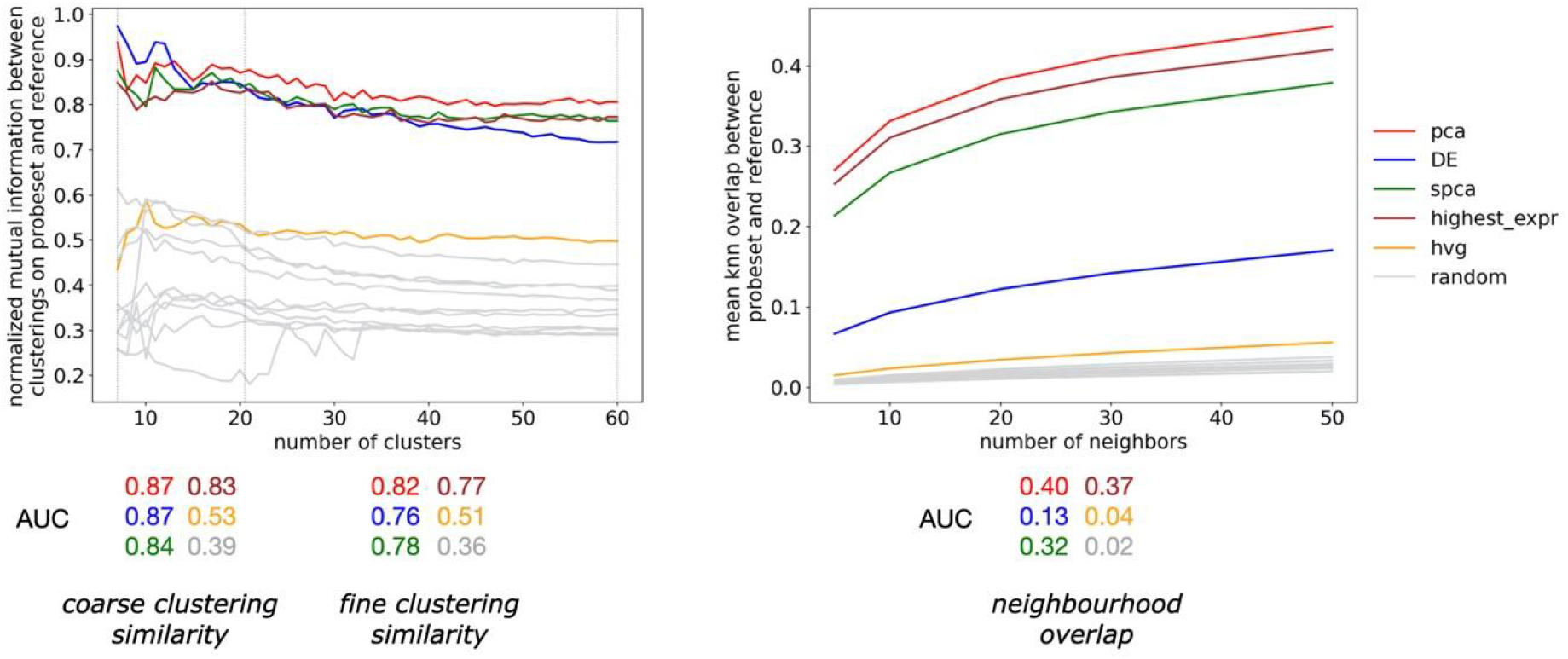
Assess variation recovery of different granularity levels via multiple metrics and parameter intervals. Clustering similarity and neighborhood overlap metrics evaluated on the Madissoon2019 dataset of gene sets with 150 genes selected with PCA, DE, SPCA, HVG, as well as highest expressed genes and random selection. The summary metrics *coarse* and *fine clustering similarity* are the AUCs of the normalized mutual information in the intervals [6,20] and [21,60] respectively, and *neighborhood overlap* is the AUC of knn overlaps over multiple k’s.

**Fig. S3:**
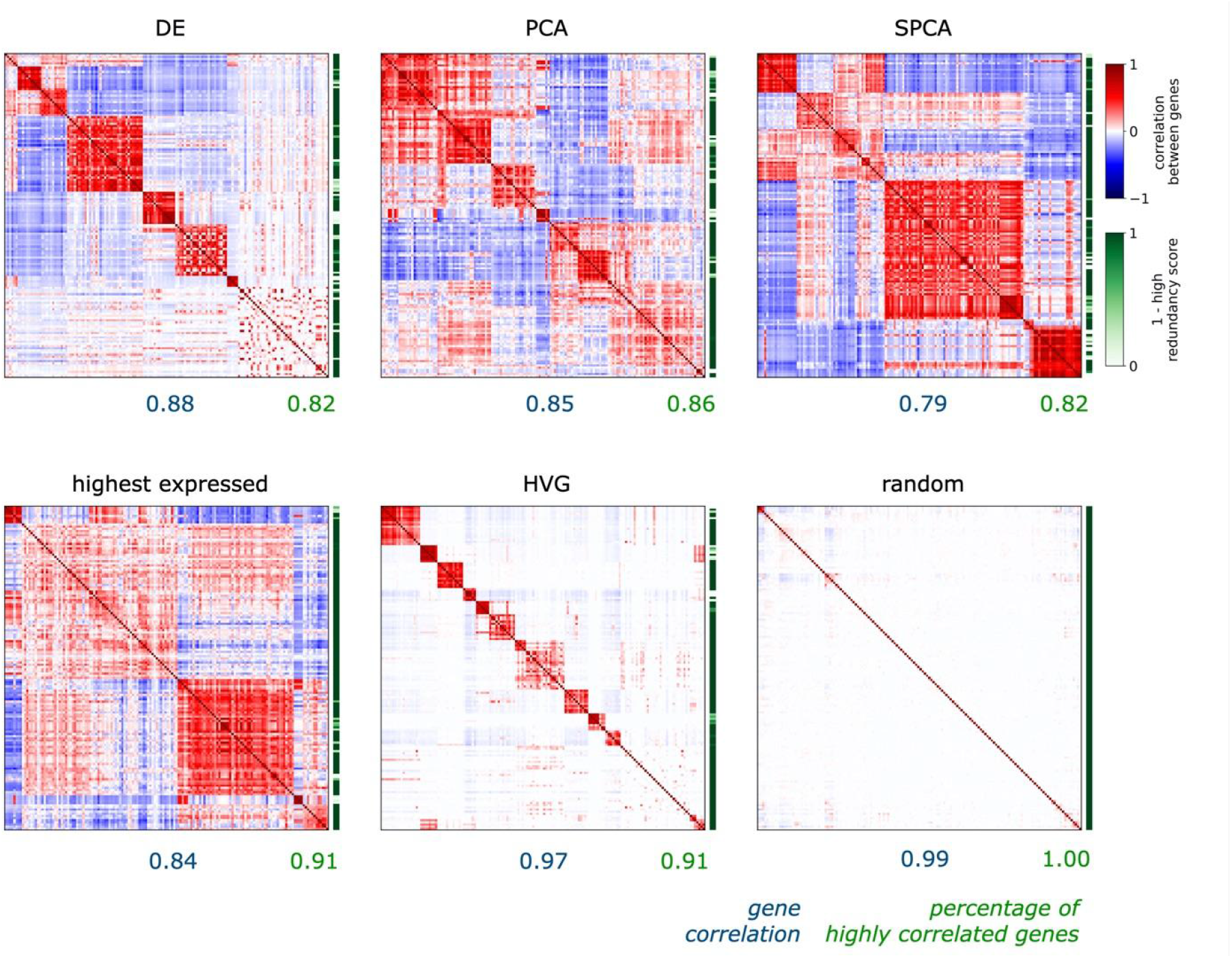
Assess gene redundancy and modular diversity via correlation evaluations. Gene correlation on the Madissoon2019 dataset of gene sets with 150 genes selected with DE, PCA, SPCA, HVG, as well as highest expressed genes and random selection. The redundancy score is a linearly smoothed step function at 0.8 of the maximal correlation of each gene. The summary metrics *gene correlation* and *percentage of highly expressed genes* are the AUCs of the normalized mutual information in the intervals [6,20] and [21,60] respectively, and *neighborhood overlap* is the AUC of knn overlaps over multiple k’s.

**Fig. S4:**
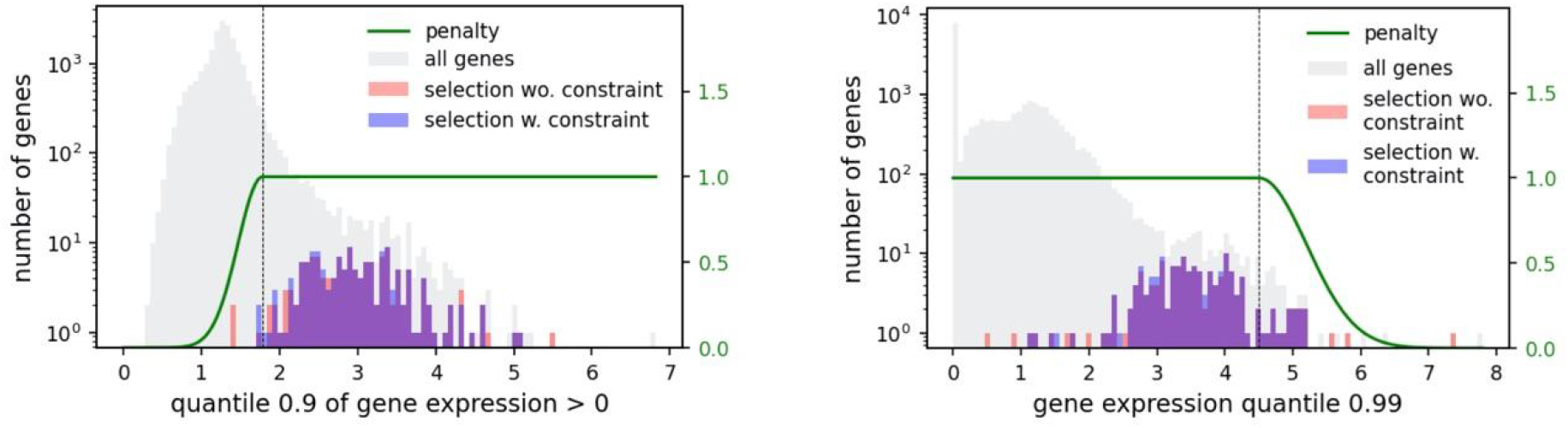
Expression constraint penalties filter out genes beyond expression thresholds. Spapros selections of 150 genes with and without expression constraint penalties. Genes are penalized and not selected as the penalties go to zero.

**Fig. S5:**
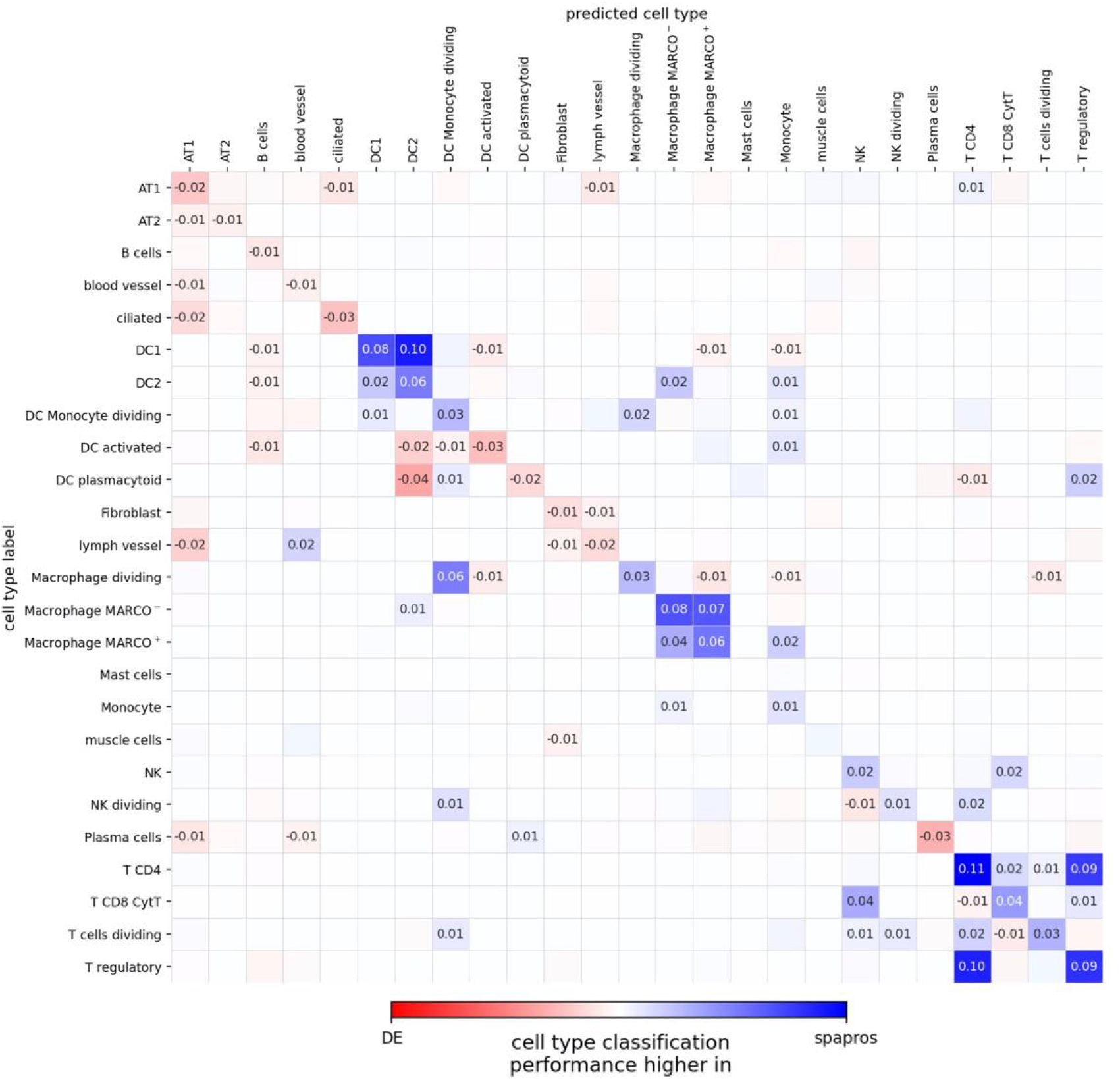
Compare cell type specific classification performance between Spapros and DE. Difference of normalized cell type classification confusion matrices between Spapros and DE selections of gene sets with 50 genes for all cell types in the Madissoon2019 dataset.

**Fig. S6:**
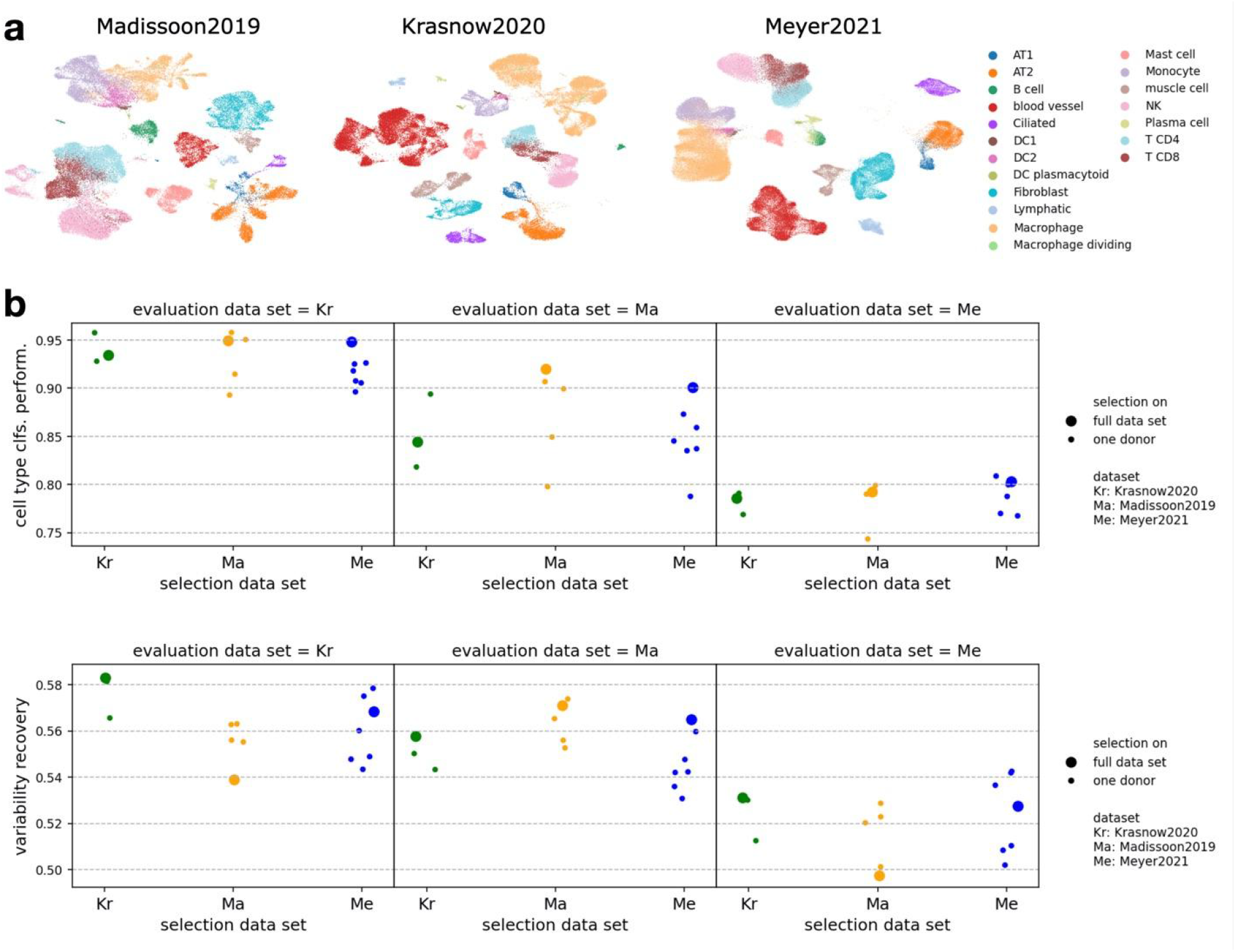
Spapros selections show robust cross dataset performance. **a**, UMAPs of the three lung datasets with unified cell type annotations for cross dataset evaluation. **b**, Cross dataset evaluations of selections on the lung data sets and on the donor samples within each data set. *Cell type clfs. perform*. is the average of the metrics *cell type classification accuracy* and *percentage of captured cell types*. Variability recovery is the average of the metrics *coarse* and *fine clustering similarity*, and *neighborhood overlap*.

**Fig. S7:**
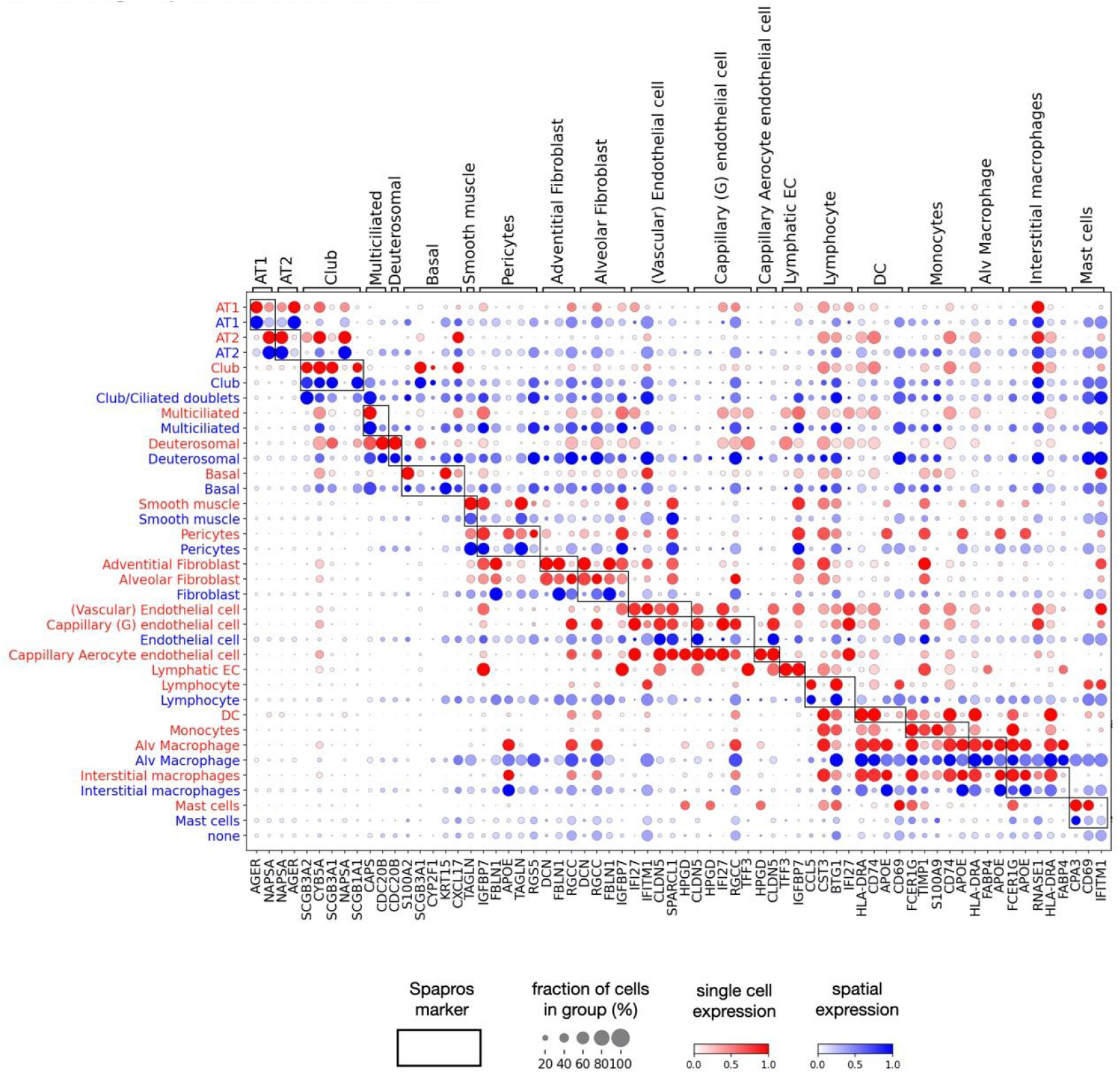
Spatial and scRNA-seq cell type clusters show similar expression profiles. Mean expressions in all spatial cell types of an intralobar lung sample (blue) and in all cell types of the single cell reference (red). The shown genes are identified as the most important genes for cell type identification in the Spapros selection.

**Fig. S8:**
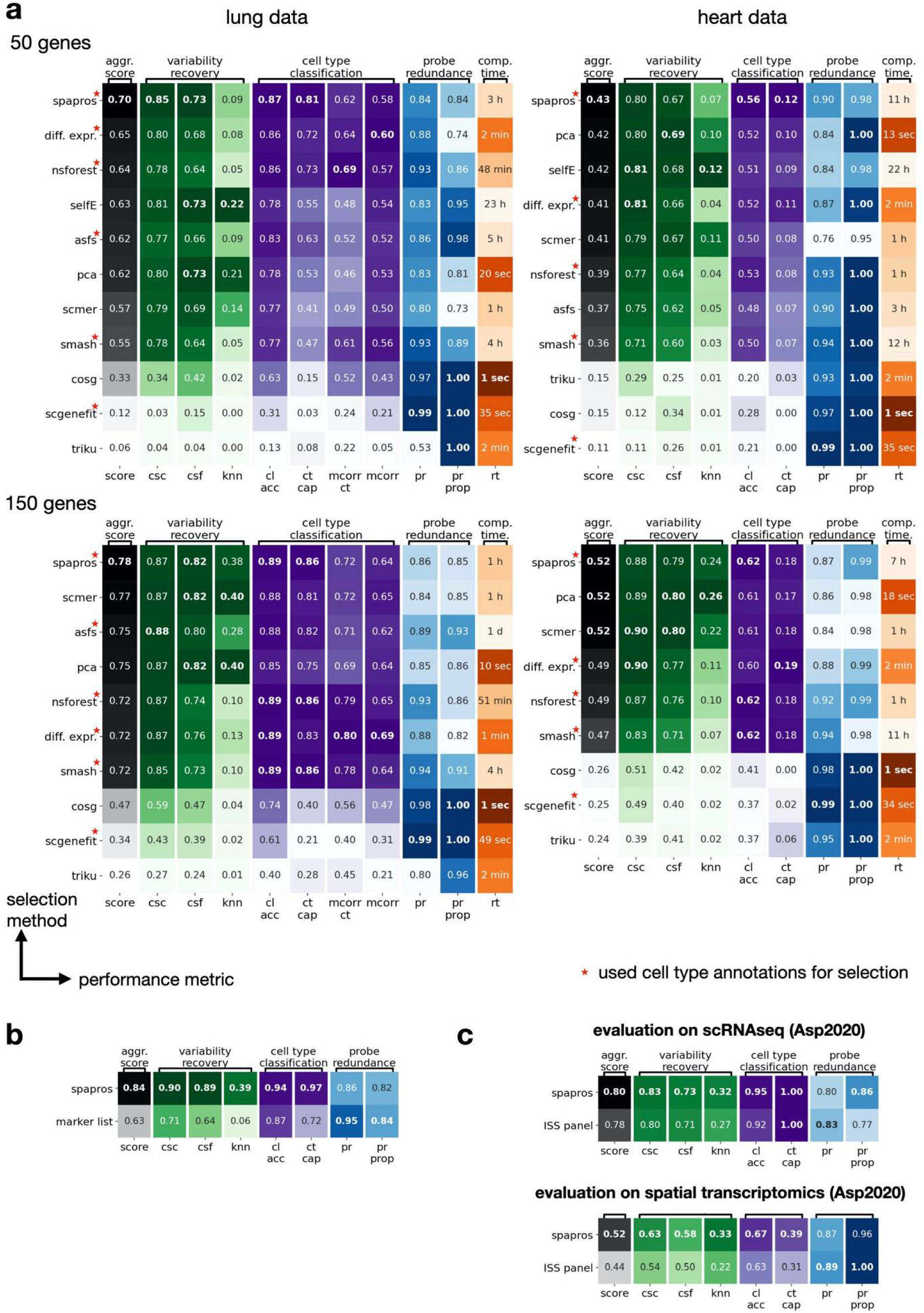
Spapros outperforms classical selection strategies and state-of-the-art methods. **a**, Heatmap of our evaluation metrics comparing Spapros with recently published methods as well as DE, and PCA-based selections. We compared selections of 50 and 150 genes for lung and heart data sets. Methods are sorted and ranked by the aggregated score of variation recovery and cell type classification. Methods that use cell type information are annotated with a red star. **b, c** Performance comparison of probe sets selected with Spapros and (**a**) a curated marker list for lung cell types and (**b**) a gene set used in ISS experiments of Asp 2020 on heart tissue which was based on selections of a single cell dataset and an untargeted spatial transcriptomics data set. Note that the cell type classification metrics on the ST dataset refer to spot based clusters instead of cell type clusters.

