## Supplementary figures and images for "Probe set selection for targeted spatial transcriptomics"

### F1_cell_outlines_intra_lobar.tif

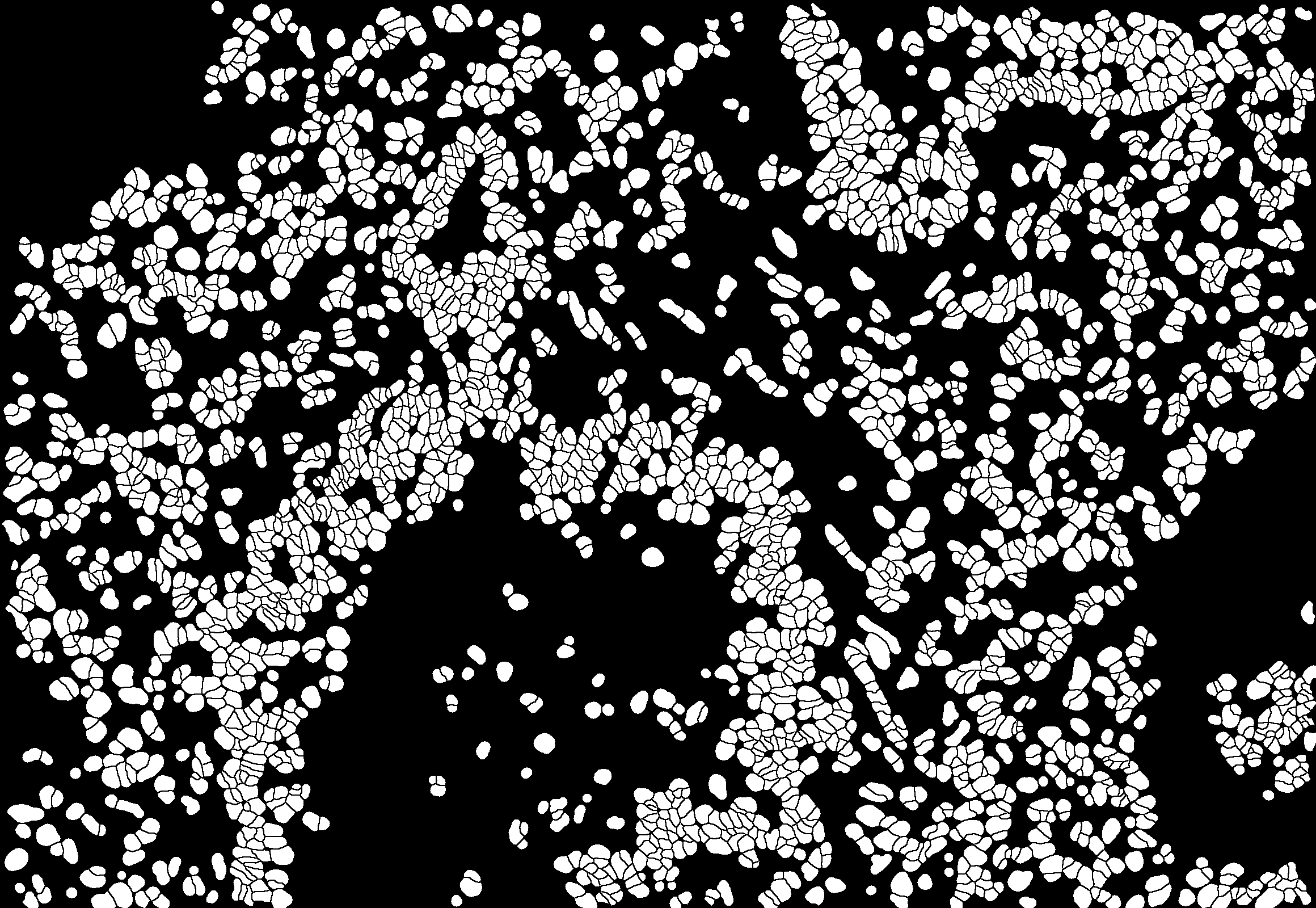
